# The transcriptomic landscapes of diverse rice cultivars grown under mild drought conditions

**DOI:** 10.1101/2020.12.11.421685

**Authors:** Taiji Kawakatsu, Shota Teramoto, Satoko Takayasu, Natsuko Maruyama, Ryo Nishijima, Yuka Kitomi, Yusaku Uga

**Affiliations:** Institute of Agrobiological Sciences, National Agriculture & Food Research Organization, Tsukuba, Ibaraki 305-8604, Japan; Institute of Crop Sciences, National Agriculture & Food Research Organization, Tsukuba, Ibaraki, 305-8602, Japan

## Abstract

Root system architecture affects plant drought tolerance and other key agronomic traits such as lodging. However, although phenotypic and genomic variation has been extensively analyzed, few field studies have integrated phenotypic and transcriptomic information, especially for below-ground traits such as root system architecture. Here, we report the phenotypic and transcriptomic landscape of 61 rice (*Oryza sativa*) accessions with highly diverse below-ground traits grown in an upland field under mild drought stress. We found that four principal components explained the phenotypic variation and that accessions could be classified into four admixture groups (admixed, *aus*, *indica*, and *japonica*) based on their tiller numbers and crown root diameters. Transcriptome analysis revealed that differentially expressed genes associated with specific admixture groups were enriched with stress response-related genes, suggesting that admixture groups have distinct stress response mechanisms. Root growth was negatively correlated with auxin-inducible genes, suggesting an association between auxin signaling and mild drought stress. A negative correlation between crown root diameter and stress response-related genes suggested that thicker crown root diameter is associated with resistance to mild drought stress. Finally, co-expression network analysis implemented with DNA affinity purification followed by sequencing (DAP-seq) analysis identified phytohormone signaling networks and key transcription factors negatively regulating crown root diameter. Our datasets provide a useful resource for understanding the genomic and transcriptomic basis of phenotypic variation under mild drought stress.

**ONE-SENTENCE SUMMARY:** Catalog of the phenomes and transcriptomes of rice cultivars grown in upland fields provides a resource for further studies toward breeding climate-resilient crops.

## INTRODUCTION

Breeding high-yielding and stress-tolerant varieties of rice (*Oryza sativa*) is crucial for sustainable food production. Root system architecture, the spatial distribution of roots with various shapes and functions (Lynch, 1995; Smith and De Smet, 2012), is an important trait for crop yield and stress tolerance (Lynch, 1995) because nutrients and water are heterogeneously distributed in the soil and affected by soil environment and cropping system (Follett and Peterson, 1988; Franzluebbers and Hons, 1996; Salinas-Garcia et al., 2002). Root system architecture is defined by multiple parameters. For example, root diameter affects the ability of roots to penetrate the soil: thicker roots tend to penetrate hard layers (Materechera et al., 1992). Root growth angle changes the distribution of roots in the soil, resulting in different responses to drought stress and affecting the plant’s ability to gather water and nutrients (Uga et al., 2013). On the one hand, roots near the topsoil (shallow-rooting plants) can acquire phosphorous (P) from P-deficient soils, because P is found in higher concentrations in the topsoil (Lynch and Brown, 2001). On the other hand, roots in the subsoil (deep-rooting plants) acquire water from deeper soil regions, allowing the plants to perform better under drought conditions (Ludlow and Muchow, 1990; Gowariker et al., 2009).

Root system architecture varies in different rice cultivars and understanding the genetic nature of this variation requires the integration of genomics and phenotyping. Recent studies have explored rice genomic diversity in the world-wide collections of rice (Wang et al., 2018b; Tanaka et al., 2020). In addition to differences among cultivars, rice is also cultivated in different field conditions, from paddy fields to upland fields. In upland fields, most of the water comes from rainfall and water in the field is easily lost due to good soil drainage (Bernier et al. 2008). Therefore, rice in upland fields tends to suffer from water shortages and characteristics such as having deep roots are important under such conditions (Bernier et al. 2008). Field cultivation conditions are complex and affected by various environmental stimuli, such as endogenous circadian rhythm, ambient temperature, plant age, and solar radiation (Nagano et al., 2012). Indeed, gene expression profiles different between laboratory and field conditions (Song et al., 2018), highlighting the importance of studying transcriptomes in field conditions. However, field phenotyping of root system architecture remains laborious and time-consuming. Recently we developed a backhoe-assisted monolith method, enabling high-throughput root phenotyping under upland conditions (Teramoto et al., 2019). In this method, root samples are collected by driving an iron tube called a cylindrical monolith into the ground with aid of a backhoe.

Studies of the transcriptomes of diverse populations help link genotype to phenotype. Increasing numbers of population-wide transcriptome studies on rice and other crops grown in the field have been conducted (Kremling et al., 2018; Groen et al., 2020). For example, one recent study identified an allele of the flowering gene *OsMADS18* with higher expression and showed its potential utility as a drought-escape gene during breeding (Groen et al., 2020). These population-wide transcriptome studies revealed the association between gene expression and traits; however, studies in the field have generally been limited to the transcriptomes of above-ground tissues (i.e. leaves), because of the difficulty in sampling below-ground tissues in the field (Yoshino et al. 2019).

Here, we compare the transcriptomes of 61 rice accessions with highly diverse root traits, grown in upland fields (Kojima et al., 2005; Uga et al., 2009; Uga et al., 2013). Phenomics analyses using our monolith method separated four admixture groups (admixed, *aus*, *indica*, and *japonica*) based on tiller numbers and crown root diameters. Transcriptome analyses revealed that each admixture group expressed distinct sets of genes for stress responses, suggesting that each admixture group has a distinct stress responsive mechanism. Our integrated analyses suggested that variation of crown root diameter in rice accessions is associated with tolerance of the mild drought conditions that occur in upland field conditions.

## RESULTS

### Natural variation of phenomes in the field

We characterized 61 rice accessions (7 admixed, 19 *aus*, 23 *indica*, and 12 *japonica*) grown in upland fields based on three above-ground traits and 15 root system architecture traits (Fig. 1, Supplemental Tables S1–3). Levels of soil water potential indicated that the upland field at the sampling stage was experiencing mild drought stress conditions for rice, which is usually grown in the paddy field (Supplemental Fig. S1). For the above-ground traits, there were no significant differences in plant height (PH) among the cultivars (Fig. 1A). By contrast, tiller number (TN) and shoot dry weight (SDW) of *japonica* cultivars were significantly lower compared to *aus* and *indica* cultivars (Figs. 1B and C). For root system architecture traits, nine root biomass-related traits (crown root length (RL_C), lateral root length (RL_L), crown root surface area (RSA_C), lateral root surface area (RSA_L), crown root volume (RV_C), lateral root volume (RV_L), number of crown root tips (NRT_C), number of lateral root tips (NRT_L), and root dry weight (RDW)) of *japonica* were significantly lower compared to *aus* and/or *indica* (Figs. 1D, E, G, H, J, K, M, N, Q). Note that we measured root system architecture traits up to 25 cm below the surface, due to the depth of the monolith. In three root system architecture topology-related traits (ratio of root length [lateral to crown root] (RL-L/C), ratio of root surface area [lateral to crown root] (RSA-L/C), and ratio of root volume [lateral to crown root] (RV-L/C)), *indica* showed higher values compared to *aus* and/or *japonica* (Fig. 1F, I, L), suggesting that *indica* allots more carbon to the production of lateral roots. For root diameter traits, crown root diameter (RD_C) of *indica* was lower compared to *aus* and/or *japonica* (Fig. 1O) whereas there were no significant differences in lateral root diameter (RD_L) (Fig. 1P). In the root distribution-related trait, there was no significant difference in ratio of deep rooting (RDR) (Fig. 1R).

**Figure 1.**
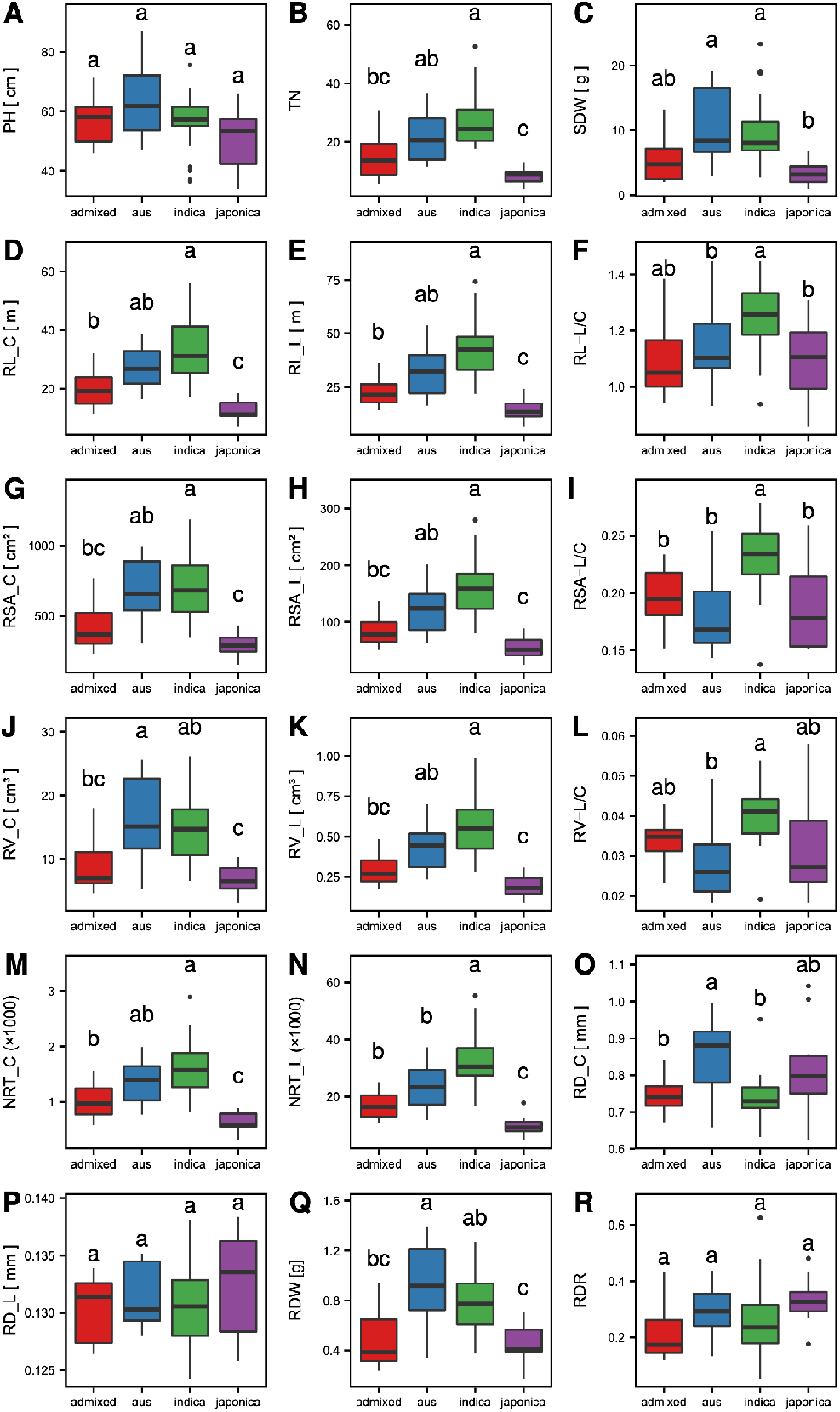
A statistical comparison among admixture groups. Four admixture groups (admixed, *aus*, *indica*, *japonica*) were compared. The graphs show three above-ground traits, plant height, PH (A), tiller number, TN (B), and shoot dry weight, SDW (C), and 15 root system architecture traits, crown root length, RL_C (D), lateral root length, RL_L (E), ratio of root length [lateral to crown root], RL-L/C (F), crown root surface area, RSA_C (G), lateral root surface area, RSA_L (H), ratio of root surface area [lateral to crown root], RSA-L/C (I), crown root volume, RV_C (J), lateral root volume, RV_L (K), ratio of root volume [lateral to crown root], RV-L/C (L), number of crown root tips, NRT_C (M), number of lateral root tips, NRT_L (N), crown root diameter, RD_C (O), lateral root diameter, RD_L (P), root dry weight, RDW (Q), and ratio of deep rooting, RDR (R). The top and bottom of the boxes mark the first and third quartiles, respectively. The center line represents the median, and the whiskers show the range of observed values within 1.5 times the interquartile range from the hinges. Values beyond, 1.5 times the interquartile range from the nearest hinge are marked by closed circles. Different letters above the plots indicate a significant difference calculated by the Steel–Dwass test.

To explore the phenotypic relationships of 18 traits in 61 rice accessions, we performed principal component (PC) analysis (Table 1). Four PCs explained 92.7% of the total phenotypic variance, and the sum of these PCs explained more than 80% of the total variation of each trait. These results indicated that the 18 traits we measured were approximately summarized by 4 PCs. We focused on traits having coefficients of over 0.40 or under −0.40, and each PC was analyzed in detail. Twelve traits were involved in PC1. Among them, nine were root-biomass related traits (NRT_C, RL_C, RSA_C, RV_C, NRT_L, RL_L, RSA_L, RV_L, RDW) and three were above-ground traits (TN, SDW, PH). This suggested that PC1 represents biomass. Six traits were involved in PC2: one above-ground trait (PH), one root diameter trait (RD_C), three root system architecture topology-related traits (RL-L/C, RSA-L/C, and RV-L/C), and one root distribution-related trait (RDR). Two traits were involved in PC3: a root diameter trait (RD_L) and the RDR trait. Only RDR was involved in PC4. Since root system architecture topology-related traits were directly affected by RD_C, and root diameter influences the root’s ability to penetrate the soil (Materechera et al., 1992), we concluded that PC2, PC3, and PC4 mainly represent crown root diameter, lateral root diameter, and root distribution in the soil, respectively.

**Table 1.** Principal component analysis of 18 traits. The coefficient values with over 0.40 or under −0.40 are in bold.

|  |  |  |  |  |  |  |  |
| --- | --- | --- | --- | --- | --- | --- | --- |
|  |  |  | coefficient |  |  |  |  |
|  | Trait | Abbreviation | PC1 | PC2 | PC3 | PC4 | % of variance |
| Above-ground | Plant height | PH | **0.62** | **-0.52** | -0.26 | -0.17 | 76.2 |
|  | Tiller number | TN | **0.91** | 0.04 | 0.07 | -0.09 | 83.6 |
|  | Shoot dry weight | SDW | **0.87** | -0.31 | -0.05 | -0.09 | 86.0 |
| Below-ground | Crown root length | RL_C | **0.98** | 0.09 | 0.10 | -0.06 | 97.8 |
|  | Lateral root length | RL_L | **0.95** | 0.27 | 0.05 | 0.05 | 98.4 |
|  | Ratio of root length (lateral to crown root) | RL-L/C | 0.35 | **0.71** | -0.29 | 0.38 | 85.5 |
|  | Crown root surface area | RSA_C | **0.98** | -0.14 | 0.04 | 0.00 | 98.3 |
|  | Lateral root surface area | RSA_L | **0.96** | 0.25 | 0.08 | 0.03 | 98.6 |
|  | Ratio of root surface area (lateral to crown root) | RSA-L/C | 0.11 | **0.96** | 0.07 | 0.06 | 95.1 |
|  | Crown root volume | RV_C | **0.92** | -0.34 | -0.02 | 0.04 | 96.6 |
|  | Lateral root volume | RV_L | **0.96** | 0.24 | 0.11 | 0.01 | 98.6 |
|  | Ratio of root volume (lateral to crown root) | RV-L/C | -0.07 | **0.96** | 0.19 | -0.06 | 96.1 |
|  | Number of crown root tips | NRT_C | **0.97** | 0.08 | 0.09 | -0.05 | 96.3 |
|  | Number of lateral root tips | NRT_L | **0.92** | 0.33 | 0.00 | 0.04 | 96.4 |
|  | Crown root diameter | RD_C | 0.20 | **-0.88** | -0.07 | 0.37 | 90.0 |
|  | Lateral root diameter | RD_L | -0.04 | -0.31 | **0.82** | 0.31 | 86.3 |
|  | Root dry weight | RDW | **0.88** | -0.39 | -0.05 | 0.07 | 94.0 |
|  | Ratio of deep rooting | RDR | -0.07 | **-0.42** | **0.58** | **0.66** | 95.6 |
|  |  | eigen value | 10.0 | 4.4 | 1.3 | 0.8 |  |
|  |  | % of variance | 56.8 | 24.3 | 7.2 | 4.4 |  |
|  |  | Cumulative % | 56.8 | 81.1 | 88.3 | 92.7 |  |

A statistical admixture group (admixed, *aus*, *indica*, *japonica*) comparison of four PC scores is shown in Fig. 2A–D. The *japonica* cultivars had a significantly lower PC1 scores (Fig. 2A) whereas *indica* had a significantly higher PC2 score (Fig. 2B). The coefficient of PC1 and root-biomass related traits was positive and that of PC2 and RD_C was negative (Table 1), suggesting that *japonica* had less root biomass in the plowed soil layer and the crown root diameter of *indica* was thinner than that of the other admixture groups. For both PC3 and PC4, there were no significant differences among *aus*, *indica*, and *japonica* (Fig. 2C, D). This implied that lateral root diameter and root distribution were influenced by local adaptation rather than admixture group divergence. Among the 12 PC1 constituents, 11 are influenced by TN because leaves and roots emerge from basal stem nodes in the main culm and tillers (Hochholdinger et al., 2004; Itoh et al., 2005). Therefore, we concluded that *aus*, *indica*, and *japonica* are classified based on two characters: tiller number and crown root diameter, which was supported by the different slopes of the scatter plots of TN and RD_C (Fig. 2E).

**Figure 2.**
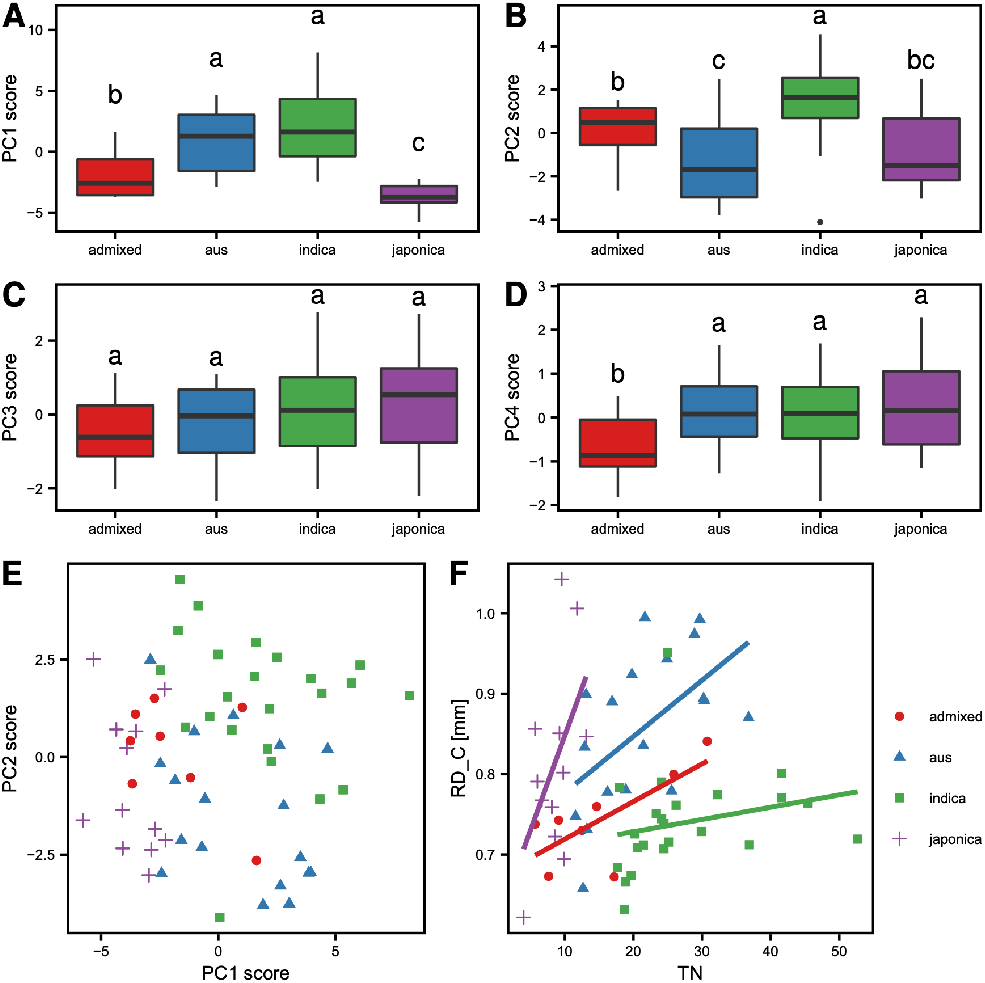
Characterization of the four admixture groups (admixed, *aus*, *indica*, *japonica*). Four principal component (PC) scores, calculated by PC analysis with three above-ground and 15 root system architecture traits, PC1 score (A), PC2 score (B), PC3 score (C), and PC4 score (D) are shown. The top and bottom of the boxes mark the first and third quartiles, respectively. The center line represents the median, and the whiskers show the range of observed values within 1.5 times the interquartile range from the hinges. Values beyond 1.5 times the interquartile range from the nearest hinge are marked by closed circles. Different letters above the plots indicate a significant difference calculated by the Steel–Dwass test. (E) A scatter diagram of PC1 and PC2. (F) A scatter diagram of regression analysis between tiller number (TN) and crown root diameter (RD_C).

### Natural variation of transcriptomes in the field

We analyzed the transcriptomes of root tips and leaves from 61 accessions grown in the same upland field. We obtained 5.3 billion raw fragments (read pairs) with a mean of 43 million fragments per sample (Supplemental Fig. S2). We detected transcripts from, on average, 17,111 genes in root tips and 15,377 genes in leaves (Fig. 3A). The numbers of expressed genes varied among admixture groups. The *japonica* accessions expressed transcripts from more genes, compared to other admixture groups, possibly because we used the *japonica* genome assembly IRGSP-1.0 as a reference genome. However, we did not observe a clear relationship between read mapping stats and admixture groups, or between numbers of uniquely mapped reads and detected genes (Fig. S2).

**Figure 3.**
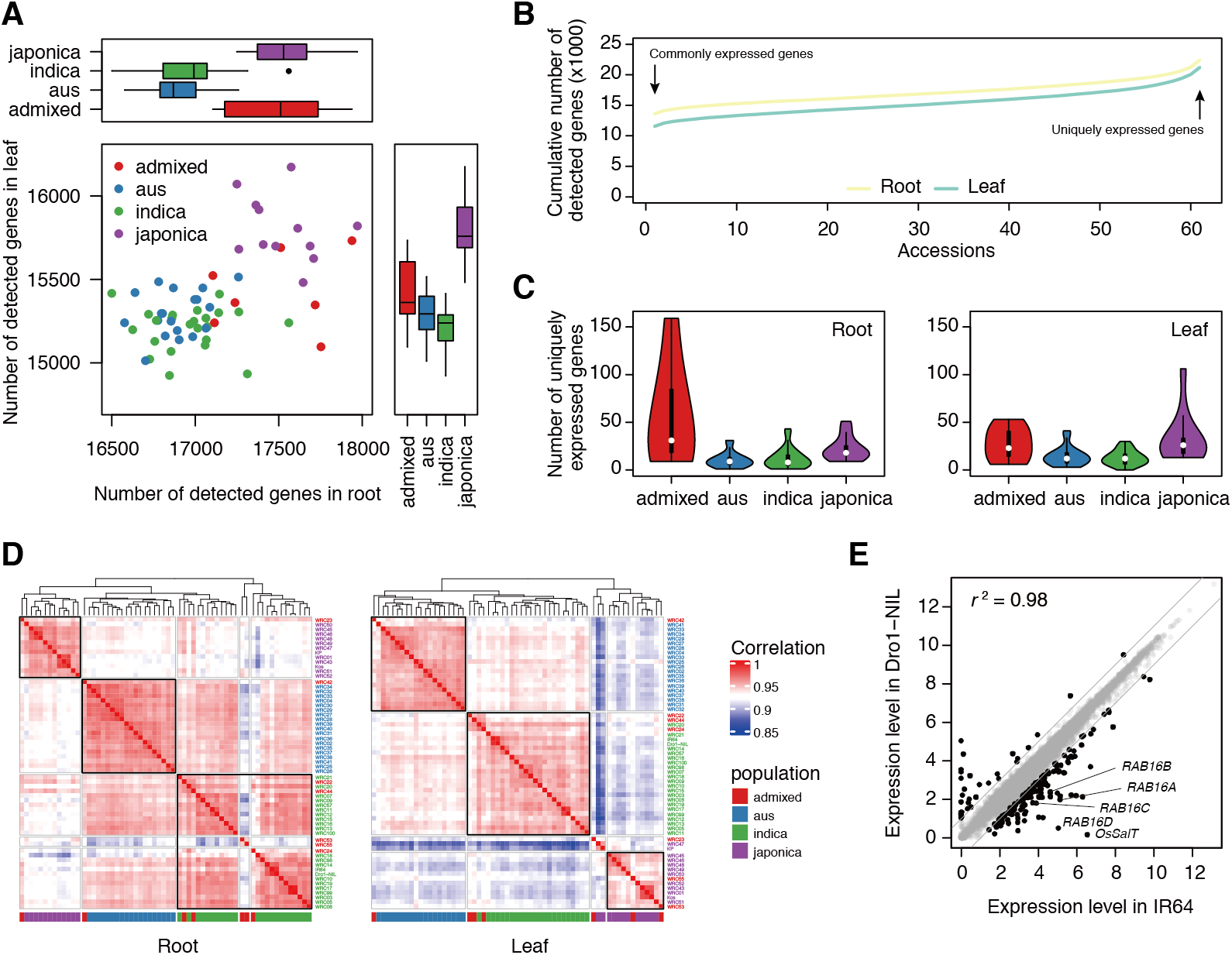
Diversity of transcriptomes in the core collection. (A) Numbers of detected genes in transcriptomes of root tips (*x*-axis) and leaves (*y*-axis). Values in the same admixture groups are summarized by boxplots. (B) Cumulative numbers of detected genes in transcriptomes of root tips and leaves. Numbers of commonly expressed genes (expressed in all accessions) and uniquely expressed genes (expressed only in one accession) are indicated. (C) Violin plots showing numbers of uniquely expressed genes in root tips (root) and leaves (leaf). (D) Heatmaps of correlation matrix of transcriptomes in root tips (left) and leaves (right). Accessions were ordered by hierarchical clustering of pairwise Pearson’s correlation coefficients. Blocks of accessions in the same admixture groups are indicated by black boxes. (E) Comparison of the expression levels of genes in root tips of IR64 and Dro1-NIL. Differentially expressed genes (|log_2_[FC]| > 1) are indicated by black points and other genes are indicated by grey points. Drought stress responsive genes are indicated.

We also compared uniquely and commonly expressed genes among groups. There were 13,633 and 11,570 commonly expressed genes in root tips and leaves, respectively, whose transcripts were detected in all accessions (Fig. 3B). By contrast, there were 1,158 and 1,164 uniquely expressed genes in root tips and leaves, respectively, whose transcripts were detected only in a single accession (Fig. 3B). Admixed accessions expressed larger numbers of uniquely expressed genes in root tips, but not in leaves (Fig. 3C). Accessions in the same admixture group had similar transcriptomes (Fig. 3D). Transcriptomes of *aus* and *indica* groups were more similar, compared to the *japonica* group, both in root tips and leaves (Fig. 3D). Transcriptomes in root tips showed higher correlations than in leaves (Fig. 3D).

Since deep rooting Dro1-NIL is genetically identical to shallow rooting IR64, except for *DRO1* locus, we compared the transcriptomes of IR64 and Dro1-NIL (Fig. 3E). Transcriptomes of IR64 and Dro1-NIL were highly correlated (r^2^ = 0.98), but there were a substantial number of differentially expressed genes (DEGs) (Fig. 3E; |log_2_[fold change (FC)]| > 1, no statistical test). These included *SALT TOLERANCE* (*OsSalT*) and *RESPONSE TO ABA* (*RAB*) genes, which are responsive to drought stress, suggesting that shallow rooting IR64 suffered from drought stress in our field conditions (Supplemental Fig. S1). Indeed, upland field cultivation could be considered a mild drought stress treatment for rice compared to paddy field conditions (Mundy and Chua, 1988; Claes et al., 1990).

Next, we identified 1,335 root tip-admixture-group-associated (root-admix) DEGs and 1,793 leaf-admixture-group-associated (leaf-admix) DEGs (|log_2_[FC]| > 1, false discovery rate [FDR] < 0.05) among admixture groups, except for the admixed group (Fig. 4). Root-admix DEGs and leaf-admix DEGs were classified into 11 clusters (R1–R11 and L1–L11, respectively), based on their level of expression (Figs. 4A and B; Supplemental Table S4). Nineteen percent (501/2627) of admixture group-associated DEGs overlapped (Fig. 4C). Shared admix-DEGs showed similar expression patterns among admixture groups, suggesting the existence of admixture group-specific gene expression patterns (Figs. 4A, B and D). For example, genes in R6 and L11 were expressed at high levels in *japonica* accessions but at low levels in *aus* and *indica* accessions. Still, there were shared admix-DEGs whose expression patterns were distinct between in root tips and leaves (e.g. genes both in R7 and L2, that were expressed at higher levels in root tips but lower levels in leaves, in *japonica* accessions) (Figs. 4A, B and D).

**Figure 4.**
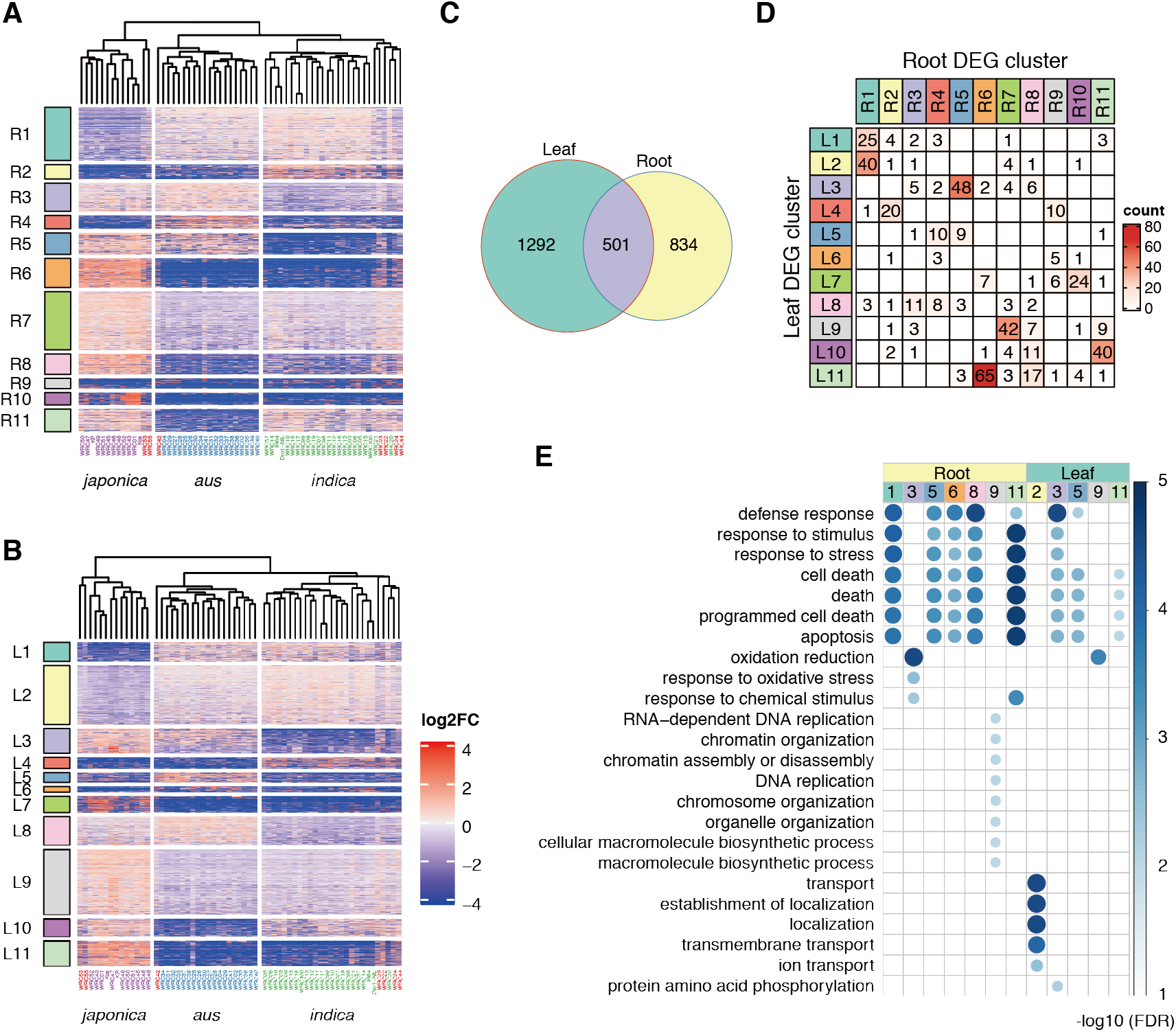
Admixture group-associated differentially expressed genes. Heatmaps of relative expression levels of admixture group-associated DEGs in root tips (A) and leaves (B). Genes were classified into 11 clusters by *k*-means clustering, based on relative expression levels. Accessions were ordered by hierarchical clustering. (C) Overlap between root tip-admixture-group-associated DEGs and leaf-admixture-group-associated DEGs. (D) Numbers of common genes between root tip-admixture-group-associated DEGs clusters and leaf-admixture-group-associated DEGs. (E) Gene ontologies enriched in each DEG cluster.

Gene ontology (GO) enrichment analysis revealed that genes for stress responses, such as defense response, response to stress, and cell death, were enriched in five and three modules in root tips and leaves, respectively (Fig. 4E), suggesting that each admixture group has distinct stress response mechanisms. Redox-related genes were enriched in R3 and expressed at lower levels in *indica* accessions than in *aus* and *japonica* accessions, whereas redox-related genes were enriched in L9 and expressed at higher levels in *japonica* accessions than in *aus* and *indica* accessions (Fig. 4A, B, E). This suggests that sensitivity and/or responsiveness to oxidative stress in root tips and leaves are associated with admixture groups.

Genes related to transport were enriched in L2 and expressed at higher levels in *indica* accessions and lower levels in *japonica* accessions (Fig. 4B, E). These genes included four sodium transporter genes: *HIGH-AFFINITY K^+^ TRANSPORTER 1;1* (*OsHKT1;1*), *OsHKT1;3*, *OsHKT2;3*, and *OsHKT2;4*. Natural variation of *OsHKT1;1* and *OsHKT1;5* affects the diversity of Na^+^ contents in roots and salt tolerance (Ren et al., 2005; Campbell et al., 2017). Na^+^ contents in root and expression levels of *OsHKT1;1* in leaves are correlated, and highest in *indica* accessions, followed in order by *aus* and *japonica* accessions. The observed differences in expression levels of these *OsHKT*s, associated with admixture groups, may partly explain the diversity of Na^+^ contents in roots among accessions (Campbell et al., 2017).

### Root traits associated with field transcriptomes in root tips

To elucidate the transcriptomic basis of root system architecture, we identified 4,370 genes whose expression patterns were correlated with root traits (|Pearson’s correlation coefficient| > 0.5). We classified root trait-associated genes into eight modules, based on correlation coefficient (Fig. 5A and Supplemental Table S5; M1–M8). Generally, the transcript levels of genes in M1, M2, M3, and M7 were negatively correlated with root growth, whereas genes in M4, M5, M6, and M8 were positively correlated with root growth (Fig. 5A).

**Figure 5.**
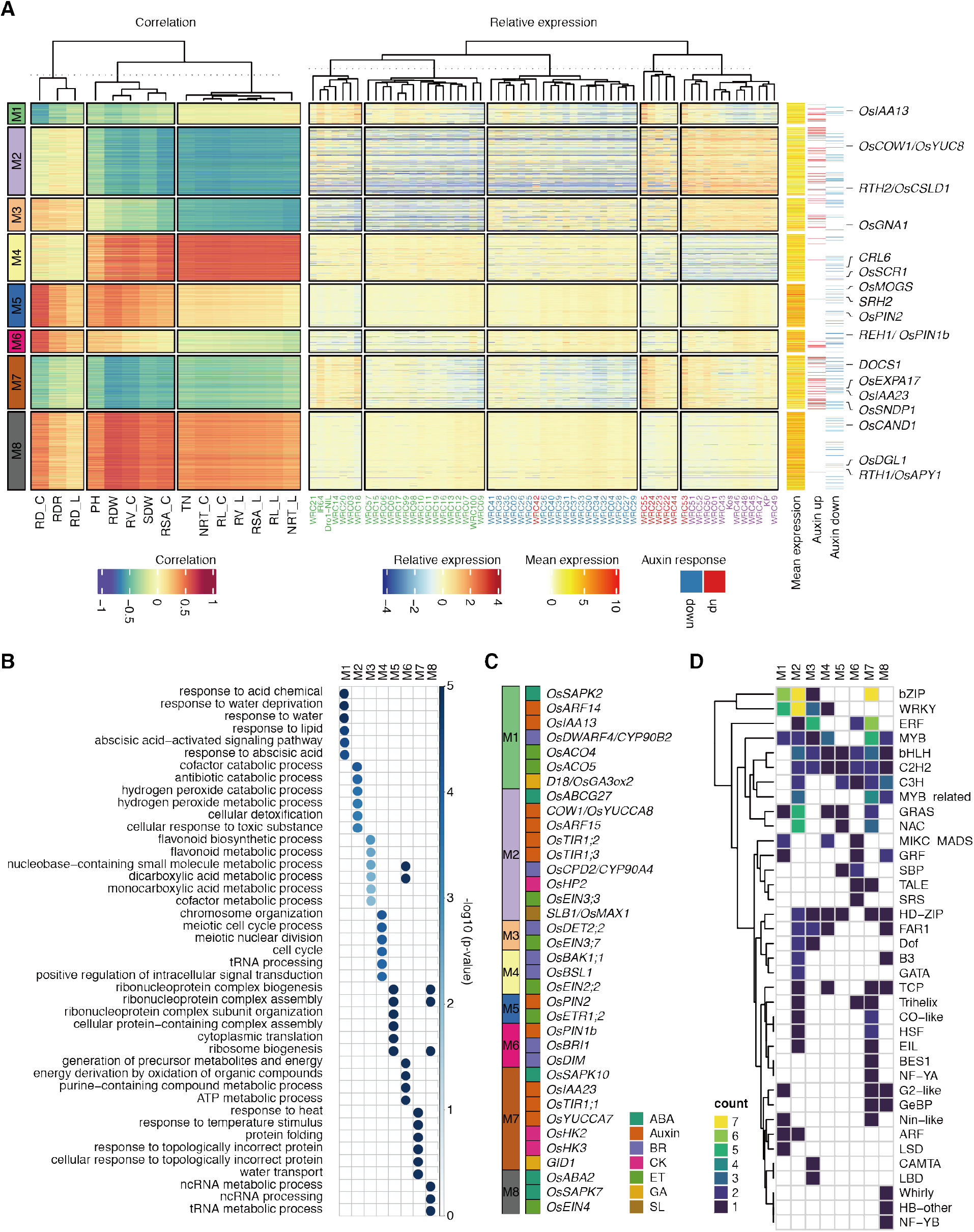
Co-expression modules in root associated with traits. (A) Heatmaps of correlations between traits and genes (left), relative gene expression (center), and others (right). Other heatmaps include mean expression of genes, auxin up-regulated genes (auxin up) and down-regulated genes (auxin down). Published root trait-associated genes are indicated at the right side of heatmaps. (B) Gene ontologies enriched in each module. The top six gene ontologies in each module are shown. (C) A list of plant hormone-related genes in each module. ABA, abscisic acid; BR, brassinosteroids; CK, cytokinins; ET, ethylene; GA, gibberellic acid; SL, strigolactones. (D) A heatmap of the numbers of transcription factor genes in each module. Transcription factor families are indicated on the right.

GO enrichment analysis revealed that genes with distinct GOs were enriched in each module (Fig. 5B). M1 was enriched with genes related to drought response, such as response to water deprivation and abscisic acid (ABA; Fig. 5B). M2 and M3 were enriched with genes related to reactive oxygen species, such as hydrogen peroxidase catabolic/metabolic process and flavonoid biosynthesis/metabolic process, respectively (Fig. 5B). M7 was enriched with genes related to heat stress, such as response to heat and protein folding (Fig. 5B). Therefore, modules negatively correlated with root growth were enriched with genes associated with abiotic stress responses. By contrast, modules positively correlated with root growth (M4, M5, M6, and M8) were enriched with energy production-associated genes and cell division genes, such as genes related to the cell cycle (M4), ribosome biogenesis (M5 and M8), generation of precursor metabolites and energy (M6) (Fig. 5B). These results reasonably explain the correlation between root growth and gene expression levels in these modules. Average gene expression levels of genes in modules negatively correlated with root growth were much lower than those in modules positively correlated with root growth, suggesting that genes in the former modules were induced by abiotic stress (Fig. 5A).

Plant hormones are involved in the formation of root system architecture. Thirty-six genes related to plant hormone biosynthesis and signaling were associated with root traits (Fig. 5C). Components of auxin signaling, the *TRANSPORT INHIBITOR RESPONSE 1* (*OsTIR1;1*, *OsTIR1;2* and *OsTIR1;3*) auxin receptor genes, and two families of transcription factor genes, i.e. the *AUXIN*/*INDOLE-3-ACETIC ACID* (*OsIAA13*, *OsIAA23*) and *AUXIN RESPONSE FACTOR* (*OsARF14* and *OsARF15*) families, belonged to modules negatively correlated with root growth (Fig. 5A). Indeed, genes up-regulated by auxin in root tips were enriched in these modules (Fig. 5A; one-sided Fisher’s exact test *p* value = 1.68e-36). Interestingly, genes down-regulated by auxin were not enriched or depleted in these modules (Fig. 5A; two-sided Fisher’s exact test *p* value = 0.30). Therefore, auxin-inducible genes were negatively associated with root growth.

Since nine root-biomass related traits were highly correlated with tiller number, we focused on root diameter, which was not affected by tiller number. RD_C was roughly clustered by admixture group (Fig. 1O). Averages of RD_C by admixture group showed a similar tendency as the root-admix DEG R3 cluster, suggesting that higher redox activity is involved in thickening of RD_C (Figs. 4A and E). Expression of genes in M1 was negatively correlated with RD_C, whereas those in M5 and M6 were positively correlated with RD_C (Fig. 5A). M1 included two genes encoding 1-aminocyclopropane-1-carboxylic acid oxidases (*OsACO4* and *OsACO5*), involved in ethylene biosynthesis, as well as the auxin signaling component genes *OsIAA13* and *OsARF14* (Fig. 5C).

Ethylene induces aerenchyma formation in the cortex, and this process depends on AUX/IAA-mediated auxin signaling (Yamauchi et al., 2020). Cortex thickness is a major determinant of root diameter (Klein et al., 2020). Hence, aerenchyma formation may be involved in reduced thickening of cortex layers, leading to thinner roots. M5 and M6 included the auxin transporter genes *OsPIN-FORMED 2* (*OsPIN2*) and *OsPIN1b*, respectively (Fig. 5A). Since OsPIN1b and OsPIN2 are responsible for acropetal and basipetal auxin transport, respectively, our data suggest that proper polar auxin transport is required for root thickening (Wang et al., 2018c). The expression level of *OsPIN2* was also correlated with RDR (Pearson’s correlation coefficient = 0.50). We found the cultivar WRC100, which had the largest RDR, also expressed *OsPIN2* at high levels (Supplemental Fig. S2). Since a loss-of-function mutant of *OsPIN2* shows small RDR, higher expression of *OsPIN2* in root tips may be responsible for the large RDR in WRC100 (Wang et al., 2018a).

### An abiotic stress signaling transcriptional network shaped the module correlated with crown root diameter in the field

Transcription factors (TFs) shape gene expression patterns and indeed, each module contained several types of TFs (Fig. 5D and Supplemental Table S6). To investigate how these modules were formed, we analyzed the co-expression network in M1 correlated with crown root diameter. M1 contained 253 genes, including 21 TFs (Fig. 5D and Supplemental Table S6). We constructed the M1 co-expression network based on the correlation of expression levels (Pearson’s correlation coefficient > 0.7). The M1 co-expression network consisted of 231 nodes (genes), including 18 TFs and 7563 edges (correlation) (Fig. 6A). We found that 83% (178/213) non-TF nodes were connected with TFs, suggesting that these genes were regulated by TFs belonging to M1. The 18 TFs in the co-expression network included six bZIPs, five WRKYs, two MYB/MYB-related TFs, and other TFs (Fig. 6A). Among them, four bZIPs (encoded by *LOC_Os01g46970*/*OSBZ8*, *LOC_Os02g52780*/*OsbZIP23*, *LOC_Os06g10880*/*OsbZIP46*, and *LOC_Os08g36790*/*TRAB1*) are involved in ABA signaling, and *LOC_Os05g34050*/*OsbZIP39* encodes a key TF in the endoplasmic reticulum-stress response (Nakagawa et al., 1996; Hobo et al., 1999; Xiang et al., 2008; Yang et al., 2011; Takahashi et al., 2012).

**Figure 6.**
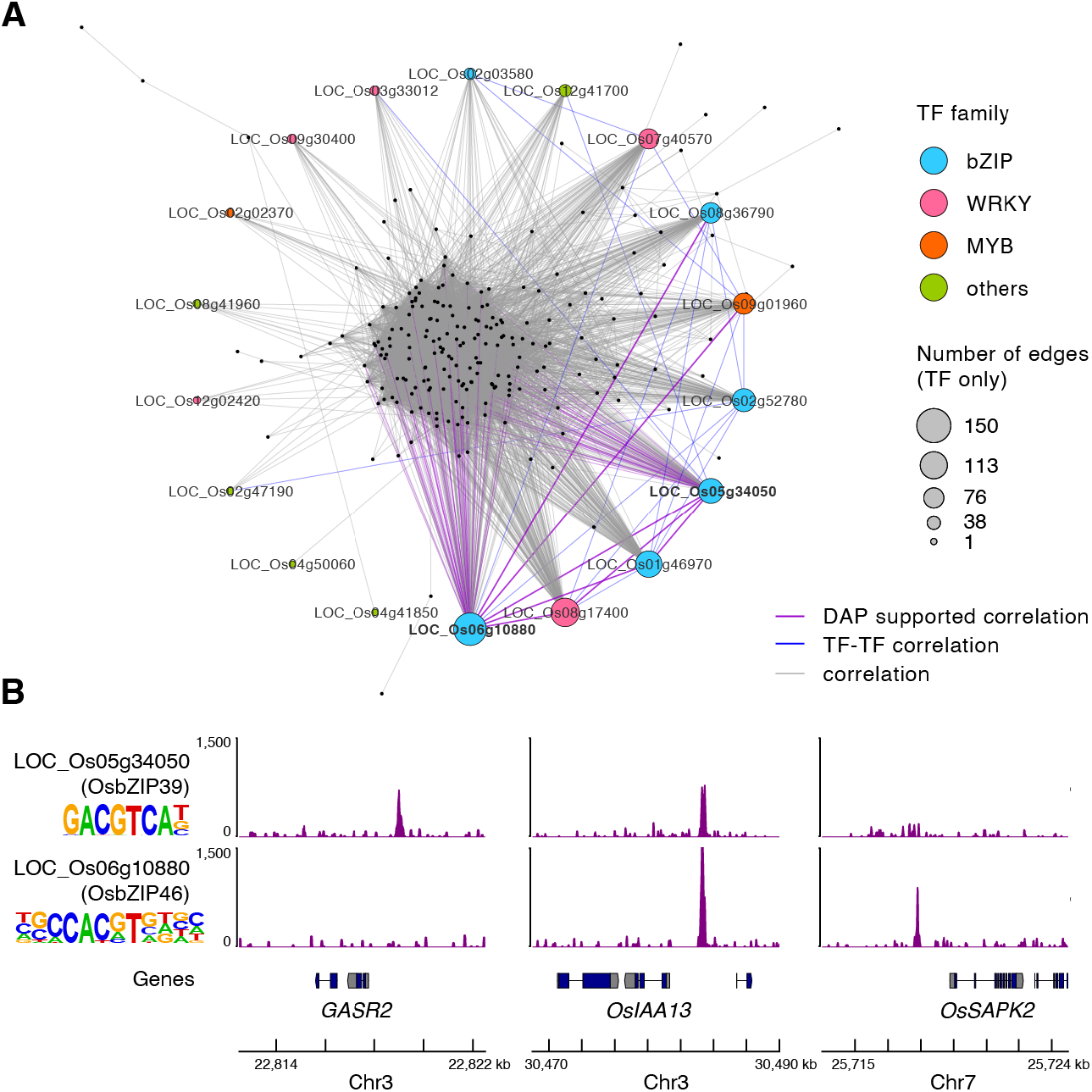
Co-expression network of the root M1 module. (A) Genes (nodes) are indicated by black dots (non-transcription factors) or circles (transcription factors, TFs). Gene pairs whose correlation coefficients were larger than 0.7 are connected by lines (edges). Circle colors represent transcription factor families. Sizes of circles represent the numbers of edges. Purple lines indicate connections that are supported by DAP-seq results. Bold purple lines are edges between transcription factors supported by DAP-seq. Blue lines are edges between transcription factors. (B) Representative DAP-seq results showing peaks upstream of plant-hormone responsive genes. Motifs enriched within the peaks are shown.

To address whether these TFs are responsible for M1 formation, we performed DNA affinity purification-followed by sequencing (DAP-seq) with N-terminally HALO-tag fused OsbZIP39 and OsbZIP46 recombinant proteins and identified 8,061 and 12,422 peaks, respectively (Supplemental Table S7) (O’Malley et al., 2016). Most enriched motifs within OsbZIP39 or OsbZIP46 bound peaks that included an ACGT core, a typical recognition motif of bZIP TFs (Fig. 6B). OsbZIP39 bound to upstream regions (within 2 kb of the transcription start site) of 24% (24/100) of nodes connected with *OsbZIP39*, whereas OsbZIP46 bound to upstream regions of 46% (64/141) of nodes connected with *OsbZIP46* (Figs. 6A). In total, 32% (69/213) non-TF genes in M1 were potential direct targets of OsbZIP39 and OsbZIP46.

Several plant hormone-related genes in M1 were targeted by OsbZIP39 and/or OsbZIP46 (Fig. 6B). For example, the gibberellin-responsive gene *GA-STIMULATED TRANSCRIPT-RELATED 2* (*GASR2*) was targeted by OsbZIP39; the auxin signaling component gene *OsIAA13* was targeted by both OsbZIP39 and OsbZIP46; the positive ABA signaling regulator *STRESS/ABA-ACTIVATED PROTEIN KINASE 2* (*OsSAPK2*) was targeted by OsbZIP46 (Kobayashi et al., 2004; Kitomi et al., 2012; Ambavaram et al., 2014). OsbZIP46 bound to the upstream regions of five TFs connected to *OsbZIP46* in M1, including *OsbZIP39*, suggesting that the predicted gene regulatory network, in which OsbZIP46 acts as a key hub TF, shapes the M1 co-expression network (Fig. 6A).

### Root traits associated with transcriptomes in leaves

Next, to address whether gene expression in leaves reflects root phenotype, we compared root traits and gene expression in leaves. We identified 1,848 genes whose expression patterns were correlated with root traits (|Pearson’s correlation coefficient| > 0.5) and classified them into four modules, based on correlation coefficient (Fig. 7 and Supplemental Table S5; M1–M4). The expression levels of genes in M2 and M3 were negatively and positively correlated, respectively, with crown root diameter (Fig. 7). GO enrichment analysis revealed that genes related to mRNA splicing and protein dephosphorylation were enriched in M2. Indeed, M2 included two protein phosphatase 2C genes, *OsPP2C9* and *OsPP2C49*, which encode negative regulators of ABA signaling (Fig. 7) (Kim et al., 2014). Furthermore, M2 included *OsABA3* and *OsSAPK9*, which are involved in ABA biosynthesis and signaling, suggesting the negative feedback regulation of ABA signaling (Fig. 7) (Kobayashi et al., 2004; Hirano et al., 2008). M3 was enriched with genes related to amino acid biosynthesis and nucleotide biosynthesis. M3 contained two cytokinin signaling component genes, *HISTIDINE KINASE 1* (*OsHK1*) and type-A *RESPONSE REGULATOR 2* (*OsRR2*), as well as the cytokinin oxidase gene *OsCKX3*, suggesting that cytokinin signaling was activated in these leaves (Fig. 7) (Ito and Kurata, 2006; Tsai et al., 2012; Zhao et al., 2020). Since drought stress decreases the amount of cytokinin in shoots (Reguera et al., 2013), our data suggest that accessions expressing genes in M3 with thick roots did not suffer from drought stress under our field conditions.

**Figure 7.**
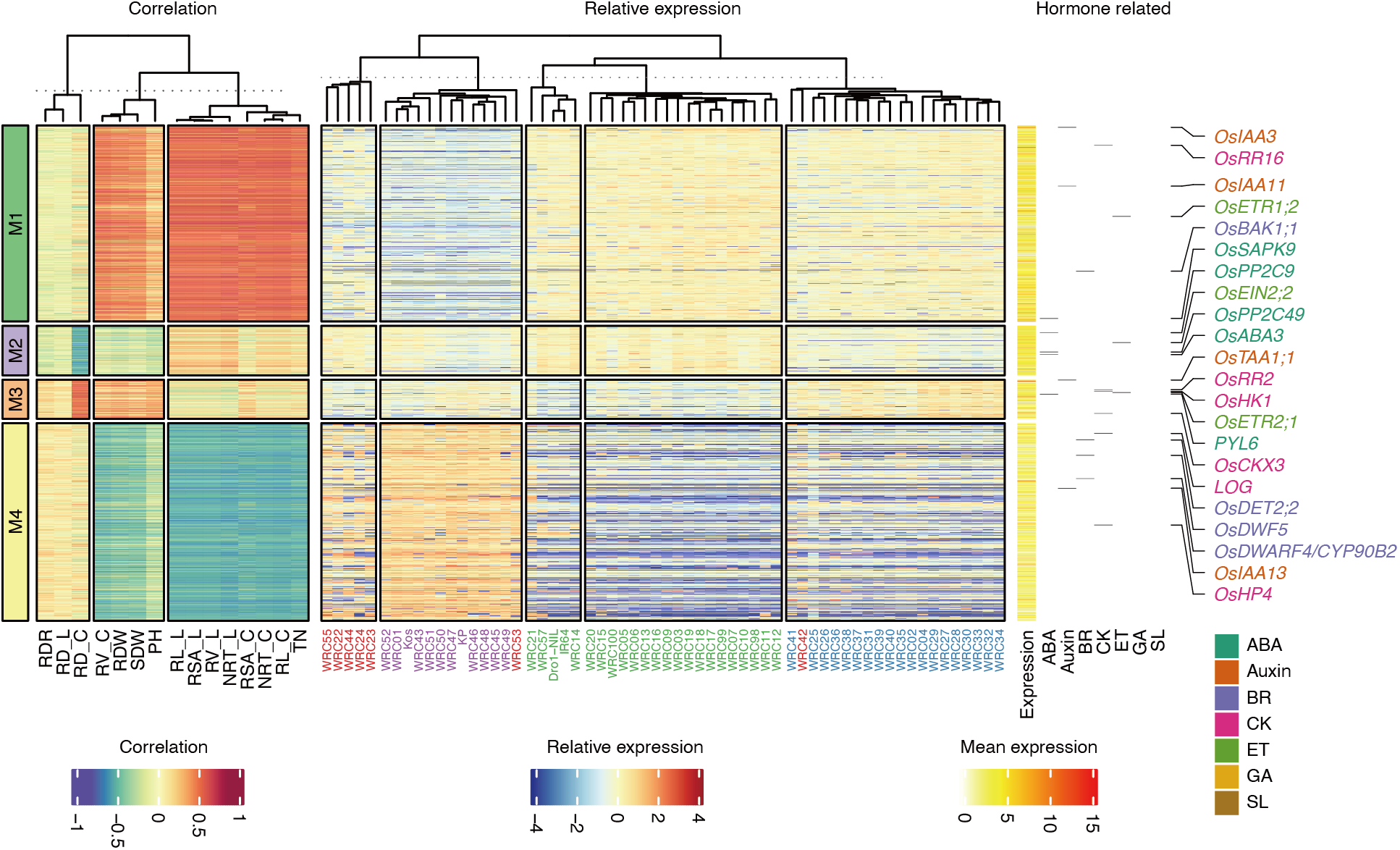
Co-expression modules in leaf associated with traits. Heatmaps of correlations between traits and genes (left), relative gene expression (center), and others (right). Other heatmaps include mean expression of genes and plant hormone-related genes. Plant hormone-related genes are indicated at the right side of the heatmaps and color coded to show the relevant hormones: ABA, abscisic acid; BR, brassinosteroids; CK, cytokinins; ET, ethylene; GA, gibberellic acid; SL, strigolactones.

## DISCUSSION

We profiled the diversity of root and shoot traits, as well as the transcriptomes of root tips and leaves in a highly diverse small population grown in upland field conditions (Kojima et al., 2005; Uga et al., 2013). The upland field conditions at sampling time were considered mild drought conditions for rice, compared to the paddy field (Fig. 3E and Supplemental Fig. S1). Integrated phenome/transcriptome analysis revealed that root traits are associated with transcriptomes of root tips and leaves. The most striking feature of the association between root traits and transcriptomes in root tips was the significant enrichment of auxin-inducible genes in co-expression modules negatively correlated with root growth (M1 in Fig. 5A), suggesting an interaction between auxin signaling and the drought stress response. Since auxin plays pivotal roles in root development, drought stress may affect auxin biosynthesis and/or signaling, leading to perturbation of auxin-mediated root developmental programs (Kitomi et al., 2018).

Co-expression network analysis and DAP-seq revealed that OsbZIP39 and OsbZIP46 directly target one-third of root M1 genes, which were negatively correlated with crown root diameter (Fig. 6). OsbZIP46 bound to five out of the top seven TFs highly connected to nodes in root M1 (Fig. 6). The expression levels of 83% of non-TF genes were correlated with those of TFs in root M1, suggesting that OsbZIP46 orchestrates a hierarchical transcriptional network in root M1. *OsbZIP46* is induced by multiple abiotic stresses and plant hormones, such as drought, heat, osmotic stress, ABA, and indole-3-acetic acid (Tang et al., 2012). OsbZIP46 is a positive regulator of ABA signaling.

Post-translational mechanisms also affect this regulatory network. OsbZIP46 physically interacts with OsSAPK2, OsSAPK6, and OsSAPK9, and is phosphorylated by OsSAPK2 and OsSAPK6 (Tang et al., 2012; Chang et al., 2017). Since phosphorylation-mimicked OsbZIP46 is constitutively active in yeast, phosphorylation by OsSAPKs probably activates OsbZIP46 (Tang et al., 2012). Indeed, overexpression of the constitutively active phosphomimic form of OsbZIP46 (but not the native form) increases tolerance to drought and osmotic stresses (Tang et al., 2012). We found that *OsSAPK2* was involved in root M1 and directly targeted by OsbZIP46, suggesting the existence of positive feedback regulation (Fig. 5C). OsbZIP23 is a close homolog of OsbZIP46 and a key player in ABA signaling in rice, but their target genes are largely different, suggesting that these bZIP TFs act independently (Tang et al., 2012). This is consistent with our DAP-seq data showing that OsbZIP46 did not directly target *OsbZIP23*.

Root M1 also included several plant hormone-related genes. For example, *OsACO4* and *OsACO5* encode ethylene biosynthesis enzymes (Fig. 5C). Although the ABA signaling pathway induces ethylene biosynthesis in Arabidopsis, the ABA signaling pathway is downstream of ethylene signaling in rice roots (Ma et al., 2014). Indeed, OsbZIP46 and OsbZIP39 did not directly target *OsACO4* or *OsACO5*, suggesting that ethylene biosynthesis is not downstream of ABA signaling; rather, enhanced ethylene production may induce ABA signaling (Fig. 6A). Rice ETHYLENE INSENSITIVE3-LIKE 1 (OsEIL1) induces the expression of *OsYUCCA8* (*OsYUC8*) and auxin biosynthesis (Qin et al., 2017). Although *OsEIL1* was not included in root co-expression modules, increased ethylene production due to upregulation of *OsACO4* and *OsACO5* may activate ethylene signaling, leading to activation of *OsYUC8*.

The auxin signaling components *OsARF14* and *OsIAA13* were included in root M1 (Kitomi et al., 2012). Local concentration gradients of auxin and subsequent auxin signaling finely control aerenchyma formation in rice roots (Yamauchi et al., 2019). ARFs are positive regulators of aerenchyma formation (Yamauchi et al., 2019). IAAs interact with ARFs and inhibit their activity; therefore, IAAs act as negative regulators of aerenchyma formation (Yamauchi et al., 2019). Although the positive regulator of *OsARF14* is unclear, *OsIAA13* was directly targeted by OsbZIP39 and OsbZIP46, suggesting that the co-expression module M1 balances aerenchyma formation (Fig. 6B).

In maize, drought-tolerant accessions have thicker roots with higher-order cortex layers and larger aerenchyma area, compared with drought-intolerant accessions, indicating that increased aerenchyma is beneficial for drought stress tolerance (Klein et al., 2020). RD_C was negatively correlated with the expression levels of ABA signaling components in root tips and leaves, suggesting that accessions with thinner roots may be more susceptible to drought conditions (Figs. 5A and 7). In maize, accessions with thicker crown roots and greater aerenchyma area perform better in drought conditions (Klein et al., 2020). Therefore, drought-stressed roots may increase aerenchyma area to adapt to drought stress with the root co-expression module M1. However, we cannot rule out the possibility that roots of drought-intolerant accessions became thinner as a stress response, because we do not have root trait data under well-watered conditions (i.e. paddy field). We also do not have information about the phenomenon of thicker crown roots and greater aerenchyma area in rice under drought conditions. Further studies would be needed to clarify these questions.

The root-admix DEG cluster R1 included the bZIP TF gene *OsFD1* (Supplemental Table S6). OsFD1 interacts with the florigen protein Heading date 3a (Hd3a) and the 14-3-3 protein GF14b, forming a florigen activation complex (Taoka et al., 2011). This complex activates *OsMADS15* in the shoot apical meristem, leading to flowering. We did not detect *OsFD1* transcripts in root tips in *japonica* accessions, but *OsFD1* was expressed in *aus* and *indica* accessions (Fig. 4A). Furthermore, *OsFD3*, a homolog of *OsFD1*, belonged to R1 (Tsuji et al., 2013). Although the functions of rice FD-like proteins are unknown, the potato (*Solanum tuberosum*) FD homologues *StFDL1a/b* are involved in tuberization (Teo et al., 2017). Our data suggest that the flowering activator OsFD1 and its homolog OsFD3 may function in root tips in *aus* and *indica* accessions, but not in *japonica* accessions. Therefore, our dataset may provide new insight into novel functions of TFs, although further study will be needed to validate these potential functions.

Using our dataset of RNA-seq reads, we identified a single nucleotide deletion in *OsSCR1* in WRC50, leading to a premature stop codon (Supplemental Fig. S4A). Arabidopsis SCARECROW (SCR) is a GRAS type TF that regulates asymmetric cell division in root meristem and leaves (Di Laurenzio et al., 1996). *OsSCR1* is also required for proper stomatal development in rice (Wu et al., 2019). Wu et al. (2019) reported that mutants of its close paralogue *OsSCR2* show normal stomatal development but *osscr1 osscr2* double mutants have severe defects in stomatal development compared to *osscr1* single mutants, suggesting that *OsSCR2* makes a minor contribution to stomatal development. However, we did not observe any defect in stomatal development in the WRC50 cultivar, which has an *osscr1* allele, suggesting that WRC50 has another TF that acts redundantly with OsSCR1 (Fig. S4D). Expression levels of *OsSCR1* and *OsSCR2* in WRC50 were similar to those in other *japonica* accessions (Figs. S4B and C). Still, we cannot exclude the possibility that OsSCR2 can substitute for OsSCR1 in WRC50.

## CONCLUSION

Our study provides a valuable dataset of above- and below-ground transcriptomes as well as phenomics data for highly diverse rice accessions grown in upland field conditions. Our dataset includes substantial variation in phenotypes and transcriptomes. Since this study was conducted under mild drought conditions, variations in phenotypes and transcriptomes are likely derived from the diversity of tolerances to drought stress as well as from the diversity of non-stress conditions. In particular, the expression levels of abiotic stress-related genes and auxin-inducible genes were correlated with root growth inhibition, likely reflecting tolerance to drought stress. The data presented in this study provide a resource for further studies toward developing climate-resilient crops.

## MATERIALS AND METHODS

### Plant materials and field conditions

Sixty-one rice accessions consisting of 57 accessions from the NIAS Global Rice Core Collection, and the cultivars Koshihikari, IR64, Kinandang Patong (KP), and Dro1-NIL were used in this study (Kojima et al., 2005; Uga et al., 2013). Koshihikari is a Japanese temperate *japonica*, the leading cultivar in Japan. IR64 and KP are an *indica* and a tropical *japonica* cultivar, respectively. Dro1-NIL is a near-isogenic line harboring KP-type *DRO1* gene in the IR64 background, resulting in intermediate-rooting properties (Uga et al., 2013). These were used as representative varieties having different rooting types in our previous study (Teramoto et al., 2019). Detailed information on the 61 lines is shown in Supplemental Table S1. Subspecies were assigned according to a previous study (McCouch et al., 2016).

The experiment was conducted concurrently with a previous study (Teramoto et al., 2019). A field experiment was conducted in 2018 at an upland field of the Institute of Crop Science (National Agriculture and Food Research Organization, Ibaraki, Japan; 36°02′89″ N and 140°09′97″ E), on a volcanic ash soil of the Kanto loam type (Humic Andosol). Fertilizer (5.2 g N m^−2^, 15.4 g P_2_O_5_ m^−2^, and 5.6 g K_2_O m^−2^) was supplied before rice planting. Three experimental blocks were designed as three replications. Each block comprised 61 plots and each plot corresponded to an accession. Each plot consisted of 15 hills in a 3×5 grid. The hill spacing was 40×40cm, and thus plot size was 1.2 × 2.0 m. Three seeds were sown in each hill on June 5. On July 3 and July 4, two seedlings were removed from each hill.

Five hills in a straight line in each plot were used for the basket assay (Uga et al., 2009). Three plastic mesh baskets (15-cm diameter and 6-cm tall) were filled with soil and buried at the three middle hills. The baskets were buried on the same days we removed the seedlings. Another five hills in a straight line in each plot were used for the backhoe-assisted monolith method (Teramoto et al., 2019). The middle three hills were used for sampling. To avoid severe drought stress, water was supplied with a sprinkler before the rice leaves started to roll.

### Measurement of above-ground traits

Three individuals in each plot were subjected to above-ground trait measurements. The tiller number (TN) and plant height (PH) were measured on July 25. Above-ground samples were collected from July 26 to August 2. The collected samples were dried at 80°C for 3 days to measure their shoot dry weight (SDW).

### Measurement of root system architecture traits

Rice roots were sampled by using the backhoe-assisted monolith method (Teramoto et al., 2019) from July 26 to August 1. The monolith was a cylinder, 20 cm in diameter and 30 cm depth, attached to a backhoe. For root sampling, the monolith was set on the ground and vertically driven into the ground to a depth of 25 cm. This depth was sufficient to collect root samples from the plow layer in upland fields (Teramoto et al., 2019). The monolith was then raised up, and root samples were washed by hand and collected from the soil within the monolith.

Root samples were stored in 70% ethanol at 10°C until use. Number of root tips (NRT), Total root length (RL), total root surface area (RSA), and total root volume (RV) were measured by using WinRHIZO Pro 2017a software (Regent Instruments, Canada). Crown roots were separated by removing shoot parts and scanned with 400 dpi resolution. On WinRhizo software, all root traits were calculated with 2 diameter classes: less than 0.2 mm (lateral roots) and greater than 0.2 mm (crown roots). Average root diameter (RD) of each class was calculated as follows:

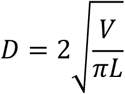

where *D* is root diameter (cm), *V* is total root volume (cm^3^), and *L* is total root length (cm). ‘_C’ and ‘_L’ were suffixed to the abbreviation to indicate the values for lateral roots or crown roots, respectively. The suffix ‘-L/C’ indicated the ratio of lateral roots to crown roots. The measured root samples were dried at 80°C for 3 days to measure their root dry weight (RDW).

Ratio of deep rooting (RDR) was measured on July 24 according to a previous study (Uga et al. 2009). The basket was dug up and the roots penetrating the basket mesh were counted. RDR was defined as the total number of crown roots penetrating the lower part of the mesh (53 to 90° to horizontal) divided by the total number of crown roots penetrating the whole mesh.

### Data processing on the phenome

RDR, TN, PH, and SDW were averaged for the plants in each plot, then were averaged again among three blocks to minimize the effect of position in the field. Traits measured by the monolith method were processed in a different way. An individual with an average tiller number was selected from each plot, and these individuals were subjected to WinRhizo analysis. Measurements were averaged across blocks. For statistical analysis, subspecies comparison was performed with a Steel–Dwass test in R (https://www.r-project.org/). A *p* value < 0.05 was considered statistically significant. Principal component (PC) analysis was performed using an R package, FactMineR (Le et al., 2008).

### RNA-seq analysis

Total RNA was extracted by the HighGI method (Yoshino et al., 2020). Equal amounts of RNA extracted from three independent plants were pooled, then RNA-seq libraries were prepared using the NEBNext Ultra II Directional mRNA-seq kit (New England Biolabs) following the manufacturer’s instructions. RNA-seq libraries were sequenced using the Illumina NovaSeq6000 at Macrogen Japan. Sequencing of libraries was performed on the S4 flow cells with paired-end 150-bp and unique dual index reads. Reads were mapped to the IRGSP-1.0 genome assembly and MU7 gene models using the STAR aligner with options “--outFilterMultimapNmax 20 --outMultimapperOrder Random --outSAMmultNmax 1 --alignIntronMax 3608 --twopassMode Basic” (Dobin et al., 2013). Uniquely mapped read counts were quantified using the featureCounts (ver. 1.6.4) with options “−s 2 −p −B −C” (Liao et al., 2014). Differentially expressed genes were called with glmQLFTest of R package edgeR (ver. 3.30.3) and fragments per kilobase exon per million reads (FPKM) values were computed after trimmed mean of M values normalization (Robinson et al., 2010). Heatmaps were visualized with the R package ComplexHeatmap (ver. 2.5.5) (Gu et al., 2016).

### Correlation between traits and gene expression levels

Pearson’s correlation coefficients between traits and gene expression levels (log_2_[FPKM + 1]) were computed. Combinations of traits and genes with absolute values of the correlation coefficient larger than 0.5 were kept and classified into clusters with *k*-means clustering. Heatmaps were visualized with the R package ComplexHeatmap (ver. 2.5.5) (Gu et al., 2016).

### DAP-seq

DAP-seq was performed as described previously, using recombinant Halo-OsbZIP39 and Halo-OsbZIP46 (O’Malley et al., 2016; Bartlett et al., 2017). Reads were mapped to the IRGSP-1.0 genome assembly using Bowtie2 (ver. 2.3.4.1) with default parameters and uniquely mapped reads were extracted using samtools (ver. 1.9) (Li et al., 2009; Langmead and Salzberg, 2012). Peaks were called using gem (ver. 2.7) with options “--k_min 6 --k_max 20 --k_seqs 600” (Guo et al., 2012). Enriched motifs within the top 1000 peaks were identified using HOMER (ver. 4.11) with default parameters (Heinz et al., 2010). Peaks were assigned to the closest genes with the closestBed command of BEDtools (ver. 2.27.1) and peaks that were more than 2 kb from any gene were discarded (Quinlan and Hall, 2010). Data were visualized with pyGenomeTracks (ver. 3.5.1) (Lopez-Delisle et al., 2020).

### Co-expression network analysis

Pairwise Pearson’s correlation coefficients between genes in the same module were computed, and gene pairs whose Pearson’s correlation coefficients over 0.7 were kept. Co-expression networks were visualized and analyzed with Cytoscape (ver. 3.8.0) (Shannon et al., 2003).

### Gene ontology analysis

GO enrichment analyses were performed with the agriGO v2.0 database (http://systemsbiology.cau.edu.cn/agriGOv2/) for admix-DEGs or with R package clusterProfiler (ver. 3.16.1) for trait-associated DEGs (Yu et al. 2012). For trait-associated DEGs, only the top six GOs in each module were shown. Data were visualized with R packages corrplot (ver. 0.85) (https://github.com/taiyun/corrplot).

## Data availability

All sequence data are available at the Gene Expression Omnibus repository (GSE162313).

## ACKNOWLEDGMENTS

We thank Hiroko Yajima, Yukie Ikemoto, Yoko Fukuda, Emiko Odajima, and the staff of the technical support section of NARO for technical support in the upland field trials. We thank the Advanced Analysis Center of NARO for the use of facilities.

---

**Supplemental Figure 1.** The environmental data in the field. Time course of soil water potential (A and B, two locations) were logged with TEROS 21 water potential sensors (METER Group, USA). The sensors were buried at a depth of 12.5 cm. Time course of air relative humidity (C) and air temperature (D) were logged with a VP-4 air sensor (METER Group, USA). All data were stored in an EM50 data logger (METER Group, USA) at 10-minute intervals.

**Supplemental Figure 2.** Statics of RNA-seq read mapping. Fractions of mapped reads in root tips, (A) and leaves (B). Accessions were sorted by fraction of uniquely mapped reads. (C) Relation between number of uniquely mapped reads (x-axis) and number of detected genes (y-axis). Circles and triangles represent root tips and leaves, respectively.

**Supplemental Figure 3.** Correlation between the expression levels of *OsPIN2* (*LOC_Os06g44970*) and RDR.

**Supplemental Figure 4.** A mutation in *OsSCR1* did not affect stomatal development. (A) A deletion detected in WRC50 RNA-seq reads (left). A dashed box is enlarged (right). Expression levels of *OsSCR1* and *OsSCR2* in root tips (B) and leaves (C). (D) Scanning electron microscopy of stomata in WRC50. Normal stomata are ordinary aligned. Bar: 100 μm.

## Notes

**FUNDING** This work was supported by JST CREST, Japan (grant number JPMJCR17O1 to Y.U. and T.K.) and JSPS KAKENHI (grant number 17H03753 to T.K.).

## LITERATURE CITED

Ahmadi N, Audebert A, Bennett MJ, Bishopp A, de Oliveira AC, Courtois B, Diedhiou A, Diévart A, Gantet P, Ghesquière A, Guiderdoni E, Henry A, Inukai Y, Kochian L, Laplaze L, Lucas M, Luu DT, Manneh B, Mo X, Muthurajan R, Périn C, Price A, Robin S, Sentenac H, Sine B, Uga Y, Véry AA, Wissuwa M, Wu P, Xu J (2014) The roots of future rice harvests. Rice (N Y) 7: 29

Ambavaram MM, Basu S, Krishnan A, Ramegowda V, Batlang U, Rahman L, Baisakh N, Pereira A (2014) Coordinated regulation of photosynthesis in rice increases yield and tolerance to environmental stress. Nat Commun 5: 5302

Atkinson JA, Rasmussen A, Traini R, Voß U, Sturrock C, Mooney SJ, Wells DM, Bennett MJ (2014) Branching out in roots: uncovering form, function, and regulation. Plant Physiol 166: 538–550

Bartlett A, O’Malley RC, Huang SC, Galli M, Nery JR, Gallavotti A, Ecker JR (2017) Mapping genome-wide transcription-factor binding sites using DAP-seq. Nat Protoc 12: 1659–1672

Bernier J, Atlin GN, Serraj R, Kumar A, Spaner D (2008) Breeding upland rice for drought resistance. J Sci Food Agric 88: 927–939

Campbell MT, Bandillo N, Al Shiblawi FRA, Sharma S, Liu K, Du Q, Schmitz AJ, Zhang C, Véry AA, Lorenz AJ, Walia H (2017) Allelic variants of OsHKT1;1 underlie the divergence between indica and japonica subspecies of rice (Oryza sativa) for root sodium content. PLoS Genet 13: e1006823

Chang Y, Nguyen BH, Xie Y, Xiao B, Tang N, Zhu W, Mou T, Xiong L (2017) Co-overexpression of the Constitutively Active Form of OsbZIP46 and ABA-Activated Protein Kinase SAPK6 Improves Drought and Temperature Stress Resistance in Rice. Front Plant Sci 8: 1102

Claes B, Dekeyser R, Villarroel R, Van den Bulcke M, Bauw G, Van Montagu M, Caplan A (1990) Characterization of a rice gene showing organ-specific expression in response to salt stress and drought. Plant Cell 2: 19–27

Di Laurenzio L, Wysocka-Diller J, Malamy JE, Pysh L, Helariutta Y, Freshour G, Hahn MG, Feldmann KA, Benfey PN (1996) The SCARECROW gene regulates an asymmetric cell division that is essential for generating the radial organization of the Arabidopsis root. Cell 86: 423–433

Dobin A, Davis CA, Schlesinger F, Drenkow J, Zaleski C, Jha S, Batut P, Chaisson M, Gingeras TR (2013) STAR: ultrafast universal RNA-seq aligner. Bioinformatics 29: 15–21

Follett R, Peterson G (1988) Surface soil nutrient distribution as affected by wheat-fallow tillage systems. Soil Science Society of America Journal 52: 141–147

Franzluebbers A, Hons F (1996) Soil-profile distribution of primary and secondary plant-available nutrients under conventional and no tillage. Soil & Tillage Research 39: 229–239

Gowariker V, Krishnamurthy V, Gowariker S, Dhanorkar M, Paranjape K (2009) The Fertilizer Encyclopedia. Wiley

Groen SC, Ćalić I, Joly-Lopez Z, Platts AE, Choi JY, Natividad M, Dorph K, Mauck WM, Bracken B, Cabral CLU, Kumar A, Torres RO, Satija R, Vergara G, Henry A, Franks SJ, Purugganan MD (2020) The strength and pattern of natural selection on gene expression in rice. Nature 578: 572–576

Gu Z, Eils R, Schlesner M (2016) Complex heatmaps reveal patterns and correlations in multidimensional genomic data. Bioinformatics 32: 2847–2849

Guo Y, Mahony S, Gifford DK (2012) High resolution genome wide binding event finding and motif discovery reveals transcription factor spatial binding constraints. PLoS Comput Biol 8: e1002638

Heinz S, Benner C, Spann N, Bertolino E, Lin YC, Laslo P, Cheng JX, Murre C, Singh H, Glass CK (2010) Simple combinations of lineage-determining transcription factors prime cis-regulatory elements required for macrophage and B cell identities. Mol Cell 38: 576–589

Hirano K, Aya K, Hobo T, Sakakibara H, Kojima M, Shim RA, Hasegawa Y, Ueguchi-Tanaka M, Matsuoka M (2008) Comprehensive transcriptome analysis of phytohormone biosynthesis and signaling genes in microspore/pollen and tapetum of rice. Plant Cell Physiol 49: 1429–1450

Hobo T, Kowyama Y, Hattori T (1999) A bZIP factor, TRAB1, interacts with VP1 and mediates abscisic acid-induced transcription. Proc Natl Acad Sci U S A 96: 15348–15353

Hochholdinger F, Park WJ, Sauer M, Woll K (2004) From weeds to crops: genetic analysis of root development in cereals. Trends Plant Sci 9: 42–48

Ito Y, Kurata N (2006) Identification and characterization of cytokinin-signalling gene families in rice. Gene 382: 57–65

Itoh J, Nonomura K, Ikeda K, Yamaki S, Inukai Y, Yamagishi H, Kitano H, Nagato Y (2005) Rice plant development: from zygote to spikelet. Plant Cell Physiol 46: 23–47

Kim H, Lee K, Hwang H, Bhatnagar N, Kim DY, Yoon IS, Byun MO, Kim ST, Jung KH, Kim BG (2014) Overexpression of PYL5 in rice enhances drought tolerance, inhibits growth, and modulates gene expression. J Exp Bot 65: 453–464

Kitomi Y, Inahashi H, Takehisa H, Sato Y, Inukai Y (2012) OsIAA13-mediated auxin signaling is involved in lateral root initiation in rice. Plant Sci 190: 116–122

Kitomi Y, Itoh J-I, Uga Y (2018) Genetic Mechanisms Involved in the Formation of Root System Architecture. *In* T Sasaki, M Ashikari, eds, Rice Genomics, Genetics and Breeding. Springer, Singapore, pp 241–274

Klein SP, Schneider HM, Perkins AC, Brown KM, Lynch JP (2020) Multiple Integrated Root Phenotypes Are Associated with Improved Drought Tolerance. Plant Physiol 183: 1011–1025

Kobayashi Y, Yamamoto S, Minami H, Kagaya Y, Hattori T (2004) Differential activation of the rice sucrose nonfermenting1-related protein kinase2 family by hyperosmotic stress and abscisic acid. Plant Cell 16: 1163–1177

Kojima Y, Ebana K, Fukuoka S, Nagamine T, Kawase M (2005) Development of an RFLP-based rice diversity research set of germplasm. Breeding Science 55: 431–440

Kremling KAG, Chen SY, Su MH, Lepak NK, Romay MC, Swarts KL, Lu F, Lorant A, Bradbury PJ, Buckler ES (2018) Dysregulation of expression correlates with rare-allele burden and fitness loss in maize. Nature 555: 520–523

Langmead B, Salzberg SL (2012) Fast gapped-read alignment with Bowtie 2. Nat Methods 9: 357–359

Le S, Josse J, Husson F (2008) FactoMineR: An R package for multivariate analysis. Journal of Statistical Software 25: 1–18

Li H, Handsaker B, Wysoker A, Fennell T, Ruan J, Homer N, Marth G, Abecasis G, Durbin R, Subgroup GPDP (2009) The Sequence Alignment/Map format and SAMtools. Bioinformatics 25: 2078–2079

Liao Y, Smyth GK, Shi W (2014) featureCounts: an efficient general purpose program for assigning sequence reads to genomic features. Bioinformatics 30: 923–930

Lopez-Delisle L, Rabbani L, Wolff J, Bhardwaj V, Backofen R, Grüning B, Ramírez F, Manke T (2020) pyGenomeTracks: reproducible plots for multivariate genomic data sets. Bioinformatics

Ludlow M, Muchow R (1990) A critical-evaluation of traits for improving crop yields in water-limited environments. Advances in Agronomy 43: 107–153

Lynch J (1995) Root Architecture and Plant Productivity. Plant Physiol 109: 7–13

Lynch JP, Brown KM (2001) Topsoil foraging—an architectural adaptation of plants to low phosphorus availability. Plant Soil 237:225–237

Ma B, Yin CC, He SJ, Lu X, Zhang WK, Lu TG, Chen SY, Zhang JS (2014) Ethylene-induced inhibition of root growth requires abscisic acid function in rice (Oryza sativa L.) seedlings. PLoS Genet 10: e1004701

Materechera S, Alston A, Kirby J, Dexter A (1992) Influence of root diameter on the penetration of seminal roots into a compacted subsoil. Plant and Soil 144: 297–303

McCouch SR, Wright MH, Tung CW, Maron LG, McNally KL, Fitzgerald M, Singh N, DeClerck G, Agosto-Perez F, Korniliev P, Greenberg AJ, Naredo ME, Mercado SM, Harrington SE, Shi Y, Branchini DA, Kuser-Falcão PR, Leung H, Ebana K, Yano M, Eizenga G, McClung A, Mezey J (2016) Open access resources for genome-wide association mapping in rice. Nat Commun 7: 10532

Mundy J, Chua NH (1988) Abscisic acid and water-stress induce the expression of a novel rice gene. EMBO J 7: 2279–2286

Nagano AJ, Sato Y, Mihara M, Antonio BA, Motoyama R, Itoh H, Nagamura Y, Izawa T (2012) Deciphering and prediction of transcriptome dynamics under fluctuating field conditions. Cell 151: 1358–1369

Nakagawa H, Ohmiya K, Hattori T (1996) A rice bZIP protein, designated OSBZ8, is rapidly induced by abscisic acid. Plant J 9: 217–227

O’Malley RC, Huang SC, Song L, Lewsey MG, Bartlett A, Nery JR, Galli M, Gallavotti A, Ecker JR (2016) Cistrome and Epicistrome Features Shape the Regulatory DNA Landscape. Cell 165: 1280–1292

Qin H, Zhang Z, Wang J, Chen X, Wei P, Huang R (2017) The activation of OsEIL1 on YUC8 transcription and auxin biosynthesis is required for ethylene-inhibited root elongation in rice early seedling development. PLoS Genet 13: e1006955

Quinlan AR, Hall IM (2010) BEDTools: a flexible suite of utilities for comparing genomic features. Bioinformatics 26: 841–842

Reguera M, Peleg Z, Abdel-Tawab YM, Tumimbang EB, Delatorre CA, Blumwald E (2013) Stress-induced cytokinin synthesis increases drought tolerance through the coordinated regulation of carbon and nitrogen assimilation in rice. Plant Physiol 163: 1609–1622

Ren ZH, Gao JP, Li LG, Cai XL, Huang W, Chao DY, Zhu MZ, Wang ZY, Luan S, Lin HX (2005) A rice quantitative trait locus for salt tolerance encodes a sodium transporter. Nat Genet 37: 1141–1146

Robinson MD, McCarthy DJ, Smyth GK (2010) edgeR: a Bioconductor package for differential expression analysis of digital gene expression data. Bioinformatics 26: 139–140

Salinas-Garcia J, Velazquez-Garcia J, Gallardo-Valdez A, Diaz-Mederos P, Caballero-Hernandez F, Tapia-Vargas L, Rosales-Robles E (2002) Tillage effects on microbial biomass and nutrient distribution in soils under rain-fed corn production in central-western Mexico. Soil & Tillage Research 66: 143–152

Shannon P, Markiel A, Ozier O, Baliga NS, Wang JT, Ramage D, Amin N, Schwikowski B, Ideker T (2003) Cytoscape: a software environment for integrated models of biomolecular interaction networks. Genome Res 13: 2498–2504

Smith S, De Smet I (2012) Root system architecture: insights from Arabidopsis and cereal crops. Philos Trans R Soc Lond B Biol Sci 367: 1441–1452

Song YH, Kubota A, Kwon MS, Covington MF, Lee N, Taagen ER, Laboy Cintrón D, Hwang DY, Akiyama R, Hodge SK, Huang H, Nguyen NH, Nusinow DA, Millar AJ, Shimizu KK, Imaizumi T (2018) Molecular basis of flowering under natural long-day conditions in Arabidopsis. Nat Plants 4: 824–835

Takahashi H, Kawakatsu T, Wakasa Y, Hayashi S, Takaiwa F (2012) A rice transmembrane bZIP transcription factor, OsbZIP39, regulates the endoplasmic reticulum stress response. Plant Cell Physiol 53: 144–153

Tanaka N, Shenton M, Kawahara Y, Kumagai M, Sakai H, Kanamori H, Yonemaru J, Fukuoka S, Sugimoto K, Ishimoto M, Wu J, Ebana K (2020) Whole-Genome Sequencing of the NARO World Rice Core Collection (WRC) as the Basis for Diversity and Association Studies. Plant Cell Physiol 61: 922–932

Tang N, Zhang H, Li X, Xiao J, Xiong L (2012) Constitutive activation of transcription factor OsbZIP46 improves drought tolerance in rice. Plant Physiol 158: 1755–1768

Taoka K, Ohki I, Tsuji H, Furuita K, Hayashi K, Yanase T, Yamaguchi M, Nakashima C, Purwestri YA, Tamaki S, Ogaki Y, Shimada C, Nakagawa A, Kojima C, Shimamoto K (2011) 14-3-3 proteins act as intracellular receptors for rice Hd3a florigen. Nature 476: 332–335

Teo CJ, Takahashi K, Shimizu K, Shimamoto K, Taoka KI (2017) Potato Tuber Induction is Regulated by Interactions Between Components of a Tuberigen Complex. Plant Cell Physiol 58: 365–374

Teramoto S, Kitomi Y, Nishijima R, Takayasu S, Maruyama N, Uga Y (2019) Backhoe-assisted monolith method for plant root phenotyping under upland conditions. Breed Sci 69: 508–513

Tsai YC, Weir NR, Hill K, Zhang W, Kim HJ, Shiu SH, Schaller GE, Kieber JJ (2012) Characterization of genes involved in cytokinin signaling and metabolism from rice. Plant Physiol 158: 1666–1684

Tsuji H, Nakamura H, Taoka K, Shimamoto K (2013) Functional diversification of FD transcription factors in rice, components of florigen activation complexes. Plant Cell Physiol 54: 385–397

Uga Y, Ebana K, Abe J, Morita S, Okuno K, Yano M (2009) Variation in root morphology and anatomy among accessions of cultivated rice (Oryza sativa L.) with different genetic backgrounds. Breeding Science 59: 87–93

Uga Y, Kitomi Y, Ishikawa S, Yano M (2015) Genetic improvement for root growth angle to enhance crop production. Breed Sci 65: 111–119

Uga Y, Sugimoto K, Ogawa S, Rane J, Ishitani M, Hara N, Kitomi Y, Inukai Y, Ono K, Kanno N, Inoue H, Takehisa H, Motoyama R, Nagamura Y, Wu J, Matsumoto T, Takai T, Okuno K, Yano M (2013) Control of root system architecture by DEEPER ROOTING 1 increases rice yield under drought conditions. Nat Genet 45: 1097–1102

Wang L, Guo M, Li Y, Ruan W, Mo X, Wu Z, Sturrock CJ, Yu H, Lu C, Peng J, Mao C (2018a) LARGE ROOT ANGLE1, encoding OsPIN2, is involved in root system architecture in rice. J Exp Bot 69: 385–397

Wang W, Mauleon R, Hu Z, Chebotarov D, Tai S, Wu Z, Li M, Zheng T, Fuentes RR, Zhang F, Mansueto L, Copetti D, Sanciangco M, Palis KC, Xu J, Sun C, Fu B, Zhang H, Gao Y, Zhao X, Shen F, Cui X, Yu H, Li Z, Chen M, Detras J, Zhou Y, Zhang X, Zhao Y, Kudrna D, Wang C, Li R, Jia B, Lu J, He X, Dong Z, Li Y, Wang M, Shi J, Li J, Zhang D, Lee S, Hu W, Poliakov A, Dubchak I, Ulat VJ, Borja FN, Mendoza JR, Ali J, Gao Q, Niu Y, Yue Z, Naredo MEB, Talag J, Wang X, Fang X, Yin Y, Glaszmann JC, Zhang J, Hamilton RS, Wing RA, Ruan J, Zhang G, Wei C, Alexandrov N, McNally KL, Leung H (2018b) Genomic variation in 3,010 diverse accessions of Asian cultivated rice. Nature 557: 43–49

Wang Y, Zhang T, Wang R, Zhao Y (2018c) Recent advances in auxin research in rice and their implications for crop improvement. J Exp Bot 69: 255–263

Wu Z, Chen L, Yu Q, Zhou W, Gou X, Li J, Hou S (2019) Multiple transcriptional factors control stomata development in rice. New Phytol 223: 220–232

Xiang Y, Tang N, Du H, Ye H, Xiong L (2008) Characterization of OsbZIP23 as a Key Player of the Basic Leucine Zipper Transcription Factor Family for Conferring Abscisic Acid Sensitivity and Salinity and Drought Tolerance in Rice. Plant Physiol 148: 1938–1952

Yamauchi T, Tanaka A, Inahashi H, Nishizawa NK, Tsutsumi N, Inukai Y, Nakazono M (2019) Fine control of aerenchyma and lateral root development through AUX/IAA- and ARF-dependent auxin signaling. Proc Natl Acad Sci U S A 116: 20770–20775

Yamauchi T, Tanaka A, Tsutsumi N, Inukai Y, Nakazono M (2020) A role for auxin in ethylene-dependent inducible aerenchyma formation in rice roots. Plants (Basel) 9

Yang X, Yang YN, Xue LJ, Zou MJ, Liu JY, Chen F, Xue HW (2011) Rice ABI5-Like1 regulates abscisic acid and auxin responses by affecting the expression of ABRE-containing genes. Plant Physiol 156: 1397–1409

Yoshino K, Nishijima R, Kawakatsu T (2020) Low-cost RNA extraction method for highly scalable transcriptome studies. Breed Sci 70: 481–486

Yoshino K, Numajiri Y, Teramoto S, Kawachi N, Tanabata T, Tanaka T, Hayashi T, Kawakatsu T, Uga Y (2019) Towards a deeper integrated multi-omics approach in the root system to develop climate-resilient rice. Molecular Breeding 39

Zhao H, Duan KX, Ma B, Yin CC, Hu Y, Tao JJ, Huang YH, Cao WQ, Chen H, Yang C, Zhang ZG, He SJ, Zhang WK, Wan XY, Lu TG, Chen SY, Zhang JS (2020) Histidine kinase MHZ1/OsHK1 interacts with ethylene receptors to regulate root growth in rice. Nat Commun 11: 518

